# IL-23 licenses the transition from preclinical autoimmunity to inflammatory arthritis through IL-17A-dependent mechanisms

**DOI:** 10.64898/2026.10.30.762860

**Authors:** Matthias Jarlborg, Alexandra Cîrciumaru, Céline Lamacchia, Laurie Vaillant, Erik Rakotoarijaonina, Koen Venken, Dirk Elewaut, Emiliana Rodriguez, Assunta Caruso, Arnaud Huard, Luca Santuari, Nylsa Chammartin, Delphine S Courvoisier, Massimo Andreatta, Santiago J Carmona, Axel Finckh, Cem Gabay

## Abstract

**Objectives:** The mechanisms driving transition from systemic autoimmunity to clinical rheumatoid arthritis (RA) remain poorly understood. We investigated the role of interleukin-23 (IL-23) in disease transition using a murine model of experimental arthritis, complemented by human transcriptomic analyses in individuals at risk of RA and RA synovial tissue.

**Methods:** Collagen-induced arthritis (CIA) was induced in less susceptible C57BL/6 mice and combined with sustained systemic IL-23 overexpression. Disease severity, histopathology, immune responses, and tissue gene expression were evaluated. The contribution of IL-17A was assessed using a neutralizing anti-IL-17A antibody. Expression of IL-23 pathway-related genes was assessed in whole-blood transcriptomic data from a large cohort of first-degree relatives of patients with RA (n=1080) and in an independent synovial RNA-sequencing dataset (n=218).

**Results:** In mice, sustained IL-23 overexpression converted a mild CIA phenotype into a highly penetrant arthritis characterized by increased expression of innate inflammatory mediators, and enhanced antigen-specific production of IL-17A and IFN-γ, despite unchanged collagen-specific antibody responses. Neutralization of IL-17A prevented arthritis development. In human whole blood longitudinal analyses, lower *RORC* and *KLRB1* expression was associated with a higher risk of RA onset. *KLRB1* expression was persistently lower in progressors, whereas *RORC* expression showed a steeper decline before diagnosis, although this difference did not remain significant after correction for multiple testing. In RA synovium, IL23R, IL23A, and CCR2 expression was increased compared with controls, while IL23R expression was higher in early than established RA.

**Conclusions:** IL-23 acts as a critical regulator of the transition from systemic autoimmunity to inflammatory arthritis by promoting IL-17A-dependent cellular and myeloid inflammatory pathways. The change in IL-23 pathway signatures in at-risk individuals and early RA supports the translational relevance of this pathway during disease initiation.

## Introduction

Rheumatoid arthritis (RA) is a chronic autoimmune disease whose pathogenesis arises from a complex interplay between genetic susceptibility and environmental exposures^1^. Increasing evidence supports the concept that RA develops along a continuum, progressing from an asymptomatic at-risk state to clinically manifest inflammatory arthritis^2^. During the earliest phases of disease, a breach of immune tolerance toward self-antigens occurs, leading to the generation of autoantibodies, most notably anti-citrullinated protein antibodies (ACPAs), a highly specific hallmark of RA^3^. Importantly, RA-associated autoantibodies can be detected years, before the onset of clinically apparent arthritis^4,5^. The appearance of RA-autoimmunity, often referred to as the “immune onset” of RA, represents a pivotal event in the disease continuum and is frequently considered the first pathogenic hit. However, only a subset of ACPA-positive individuals will ultimately develop arthritis, indicating that additional pathogenic events are required for disease progression. The immune mechanisms underlying the transition to arthritis remain incompletely understood. Growing evidence suggests that mucosal tissues, including the lung, oral cavity, and gut, may constitute key sites of tolerance breakdown and early immune activation^6^. In this context, disruption of epithelial barrier integrity and the release of inflammatory mediators from stressed mucosal tissues have been proposed as critical events promoting disease progression^6^.

According to the “multiple-hit” model of RA pathogenesis, additional inflammatory signals are required to drive the transition from preclinical autoimmunity to overt arthritis. Among these signals, interleukin-23 (IL-23) has emerged as a compelling candidate^7^. IL-23 is a pleiotropic cytokine with a prominent role at barrier tissues, where it orchestrates innate and adaptive immune responses, promotes T-cell polarization - including both αβ T cells and γδ T cells - and enhances antigen presentation and humoral immune responses^8–10^. Although IL-23 is widely recognized as a central pathogenic mediator in psoriatic arthritis, its contribution to RA appears to be context- and stage-dependent^11^. Clinical trials targeting IL-23 in established RA have yielded disappointing results, suggesting a limited role in established disease^12^. The apparent discrepancy between the failure of IL-23 blockade in RA and the evidence implicating IL-23 in disease initiation suggests a temporally restricted pathogenic role. Indeed, accumulating evidence from both human studies and experimental models indicates that IL-23 may be critically involved during the earliest phases of RA development. Elevated IL-23 levels have been reported in patients with early RA and have been associated with the formation of ectopic lymphoid structures that support local autoantibody production^13,14^. Experimental studies further demonstrated that IL-23 signaling contributes to the initiation of autoimmune arthritis^15,16,7^ and that therapeutic blockade of IL-23 ameliorates disease when administered before, but not after, the onset of clinical manifestations^17^. Despite these observations, the role of IL-23 during the preclinical stages of RA, particularly in the context of mucosal inflammation and epithelial stress, remains poorly defined.

We therefore hypothesized that IL-23 acts as a critical catalyzer of the transition from systemic autoimmunity to inflammatory arthritis. To test this hypothesis, we combined a murine model of autoimmune arthritis based on type II collagen immunization with analyses in individuals at risk and patients with early RA. Using this approach, we sought to determine whether IL-23 represents a key mechanistic driver of disease transition and a potential therapeutic target during the earliest stages of RA pathogenesis.

## Methods

### Mice and collagen-induced arthritis (CIA)

Male and female C57BL/6 mice aged 8–12 weeks were purchased from Charles River Laboratories (Wilmington, MA, USA) and housed in a conventional animal facility in individually ventilated cages (IVCs) under controlled environmental conditions with ad libitum access to food and water. All animals were acclimatized for at least 2 weeks before the start of the experiments. To induce collagen-specific autoimmunity, all mice were immunized with chicken type II collagen (CII; MD Bioproducts GmbH Zürich, Switzerland, Cat. 804002-lyo) emulsified in Freund’s adjuvant (Cat. F5506) completed (CFA) with heat-killed *Mycobacterium tuberculosis* H37Ra (Difco, Franklin Lakes, NJ, USA). The emulsion was prepared using CII at a final concentration of 4 mg/mL. Mice received 100 μL of emulsion by subcutaneous injection at the base of the tail (50 μL per side) on days 0 and 21. Systemic IL-23 overexpression was achieved by hydrodynamic tail vein delivery of extracellular expression vectors (EEVs), as previously described^7^. The experimental timeline was defined as follows: primary CII immunization on day 0, EEV administration on day 10, booster CII immunization on day 21, and sacrifice at the study endpoint. Briefly, on day 10 after primary immunization and before booster immunization, mice received 40 ng of an IL-23-expressing EEV (System Biosciences, Palo Alto, CA, USA) or the corresponding empty control vector (CIA controls).

To investigate the contribution of IL-17A signaling to disease development, a parallel cohort of mice received a neutralizing anti-IL-17A monoclonal antibody (BZN035; Novartis, Basel, Switzerland) or an isotype matched anti-Cyclin A antibody. Antibody treatment was initiated on day 9 after immunization (one day prior to the vector injection) and administered intraperitoneally every 48 hours until sacrifice on day 35. Each injection contained 150 μg of anti-IL-17A antibody or its isotype control. Clinical assessment was performed three times per week throughout the study and included body weight monitoring and evaluation of arthritis severity using a semi-quantitative visual analogue (VAS) scoring system. Each paw was scored on a scale from 0 to 3 as follows: 0, no visible signs of inflammation; 1, mild swelling and/or erythema affecting one or more digits; 2, moderate swelling involving the paw; and 3, severe swelling and deformity of the entire paw (based on reference^18^). Animals were monitored for a total of 45 days following primary immunization.

At the experimental endpoint, mice were euthanized by pentobarbital overdose followed by cervical dislocation. Blood was collected by cardiac puncture before cervical dislocation, and plasma was isolated for cytokine quantification and anti-CII antibody measurements. Spleens and draining lymph nodes (popliteal and inguinal) were harvested for *ex vivo* culture and antigen-specific restimulation assays. Hind paws were collected for histological assessment and gene expression analyses.

### Patient samples and transcriptomic data

Participants were recruited from the Swiss SCREEN-RA cohort, which includes first-degree relatives (FDRs) of patients with RA enriched for individuals at increased risk of developing RA^19^. Risk stratification was based on the presence of genetic risk factors, RA-related autoimmunity, musculoskeletal symptoms (arthralgia), and clinically evident arthritis. Participants were categorized into five groups with increasing risk of disease progression, as previously described^19^. Briefly, Group 1 comprised control participants, defined as first-degree relatives of patients with RA who had neither genetic risk factors nor evidence of RA-related autoimmunity. Group 2 included asymptomatic individuals with genetic susceptibility, defined by the presence of two shared epitope alleles, without autoimmunity. Group 3 consisted of individuals with RA-related autoimmunity, defined by seropositivity for anti-citrullinated protein antibodies (ACPA), rheumatoid factor levels exceeding three times the upper limit of normal, and/or the presence of anti-RA33 autoantibodies. Group 4 comprised individuals with clinically suspect arthralgia (CSA; score ≥4) or undifferentiated arthritis, without genetic susceptibility and/or RA-related autoimmunity. Group 5 comprised individuals with clinically suspect arthralgia or undifferentiated arthritis, and genetic susceptibility and/or RA-related autoimmunity. Progressors were participants in groups 1–5 at their first blood sample who developed RA during follow-up (group 6, n=30); all others were non-progressors. Follow-up ran from the first blood sample to the diagnosis of RA or the last follow-up visit, and only samples collected before diagnosis were analysed.

Whole-blood bulk RNA-sequencing data from SCREEN-RA participants were analyzed to assess expression of IL-23 pathway-related genes. Transcriptomic profiling was performed at the Health 2030 Genome Center (https://www.health2030genome.ch/; Campus Biotech, Geneva, Switzerland) on n=1080 RNA samples isolated from whole blood collected from RA-FDRs (n=390) enrolled in the SCREEN-RA cohort. RNA libraries were prepared using the Illumina TruSeq Stranded mRNA Library Preparation Kit according to the manufacturer’s instructions. Libraries were sequenced on Illumina NovaSeq 6000 and HiSeq 4000 platforms. Variance-stabilized transformed (VST) expression values generated using the DESeq2 workflow were analyzed for IL-23 pathway-related genes (genes of interest available in supplementary table 1). Only transcripts with a minimum of 10 read counts per sample in at least 20% of the samples were retained.

Publicly available bulk RNA-sequencing data from synovial biopsies (series GSE89408) were extracted from the Gene Expression Omnibus. The dataset comprises 218 samples from healthy controls and patients with arthralgia, osteoarthritis, undifferentiated arthritis, early RA, and established RA.

Raw read counts from whole blood and publicly available synovial tissue were process with the DESeq2 package. Fastq files were pre-processed using fastp^20^ version 1.3.3 for read quality control and adaptor trimming. Transcript abundance was estimated using salmon version 1.10.0^21^, with the GENCODE v45 annotations as the reference transcriptome, and the entire GRCh38 genome assembly as a decoy. Transcript count estimates were imported into R using tximport, and variance-stabilized transformed (VST) expression values were calculated using DESeq2 version 1.50.2^22^.

For the longitudinal SCREEN-RA analyses, only samples collected before RA diagnosis were included. Participants who had RA at their first sampling visit (n = 13) or received another diagnosis during follow-up (n = 11) were excluded. Genes below the detection limit in more than 30% of samples (within 0.5 VST units of the panel minimum) were flagged and not used in progression analyses.

### Statistical analysis

Batch effects between cohort (SYSCID and the PErPREV-CID) were corrected using the limma package^23^. Differential comparisons between groups were performed using non-parametric Kruskal–Wallis tests followed by multiple comparisons with false discovery rate (FDR) correction. To account for the non-independence of repeated samples obtained from the same patient, generalized estimating equations (GEE) were fitted using the geepack package in R (version 4.4.2). Gene expression was modeled as the dependent variable and study group as the independent variable. Patient identifier was specified as the clustering variable, and a Gaussian distribution with identity link was used. An independence working correlation structure was specified. Robust sandwich standard errors were used for statistical inference. P-values between groups were estimated using a Wilcoxon rank-sum test (*** p<0.001; ** p<0.01; * p<0.05).

Risk groups 2–6 were compared with group 1 using linear mixed-effects models (1071 samples from 387 participants), with a random intercept per participant and adjustment for age and sex; differences are reported as log2 fold changes. The association between gene expression and progression to RA was estimated with time-varying Cox models (time since inclusion), in which expression and ACPA and RF status were updated at each pre-diagnostic sample. Each of the 22 genes was modelled separately, adjusted for age, sex, ACPA and RF (HR per 1 SD), and genes were then considered jointly by stepwise selection with these covariates forced in. Effect modification by serostatus was tested with a gene × seropositivity interaction. Expression over time was compared between progressors and non-progressors with linear mixed-effects models (random intercept per participant) including time to diagnosis or last follow-up, group and their interaction, adjusted for age and sex; additional adjustment for ACPA and RF was a sensitivity analysis. p-values were corrected across the 22 genes with the Benjamini–Hochberg method, and q < 0.05 was considered significant. Analyses were performed in R 4.6.0 (survival, lme4, lmerTest, emmeans).For mouse experiments, data are presented as individual values with mean ± standard error of the mean (SEM). Comparisons between two groups were performed using unpaired Student’s t-test, while multiple group comparisons and longitudinal analyses were analyzed using two-way ANOVA for repeated measures, as appropriate. Normality of data distribution was assessed prior to statistical testing. Two-sided p-values <0.05 were considered statistically significant. Statistical significance was defined as follows: p < 0.05 (*), p < 0.01 (**), p < 0.001 (***), and p < 0.0001 (****). All analyses and graphical representations were performed using GraphPad Prism version 10 (GraphPad Software, La Jolla, CA, USA) or RStudio (R Foundation for Statistical Computing, Vienna, Austria, version 4.4.2).

### Ethics

The SCREEN-RA cohort has been approved by the relevant ethics committees (SCREEN-RA: 2025-00114, SYSCID: 2019-00707, PerPrevCID: 2026-00415). All experiments were approved by the Geneva Cantonal Authority for Animal Experimentation (licenses GE389, GE550, GE550A).

### Patient and public involvement

All participants provided written informed consent before enrolment in accordance with the Declaration of Helsinki. Patients and the public were not involved in the design, conduct, reporting, or dissemination of this research.

## Results

### IL-23 overexpression promotes arthritis development in collagen-induced C57BL/6 mice

Our previous work demonstrated that sustained systemic IL-23 overexpression induced inflammatory arthritis in HLA-DR4 transgenic mice immunized with citrullinated peptides, identifying IL-23 as a key driver of autoimmune arthritis^7^. We therefore investigated whether IL-23 functions more broadly as a determinant of disease transition by overcoming resistance to arthritis in a collagen-induced arthritis model. To this end, we combined the immunization against type II collagen with IL23EEV overexpression in C57BL/6 mice (Figure 2A), a strain known to be less susceptible to CIA induction^24^. This experimental approach was designed to model the effect of an additional inflammatory stimulus in the setting of established antigen-specific autoimmunity but limited disease susceptibility. Whereas CIA mice receiving the empty vector developed only mild and low incident articular manifestations (predominantly limited to isolated digit inflammation), IL-23OE mice exhibited severe polyarthritis with high penetrance and significant body weight loss (Figure 1B-C). Importantly, IL-23EEV administration alone did not induce arthritis in non-immunized C57BL/6 mice (supplementary figure 1).

**Figure 1.**
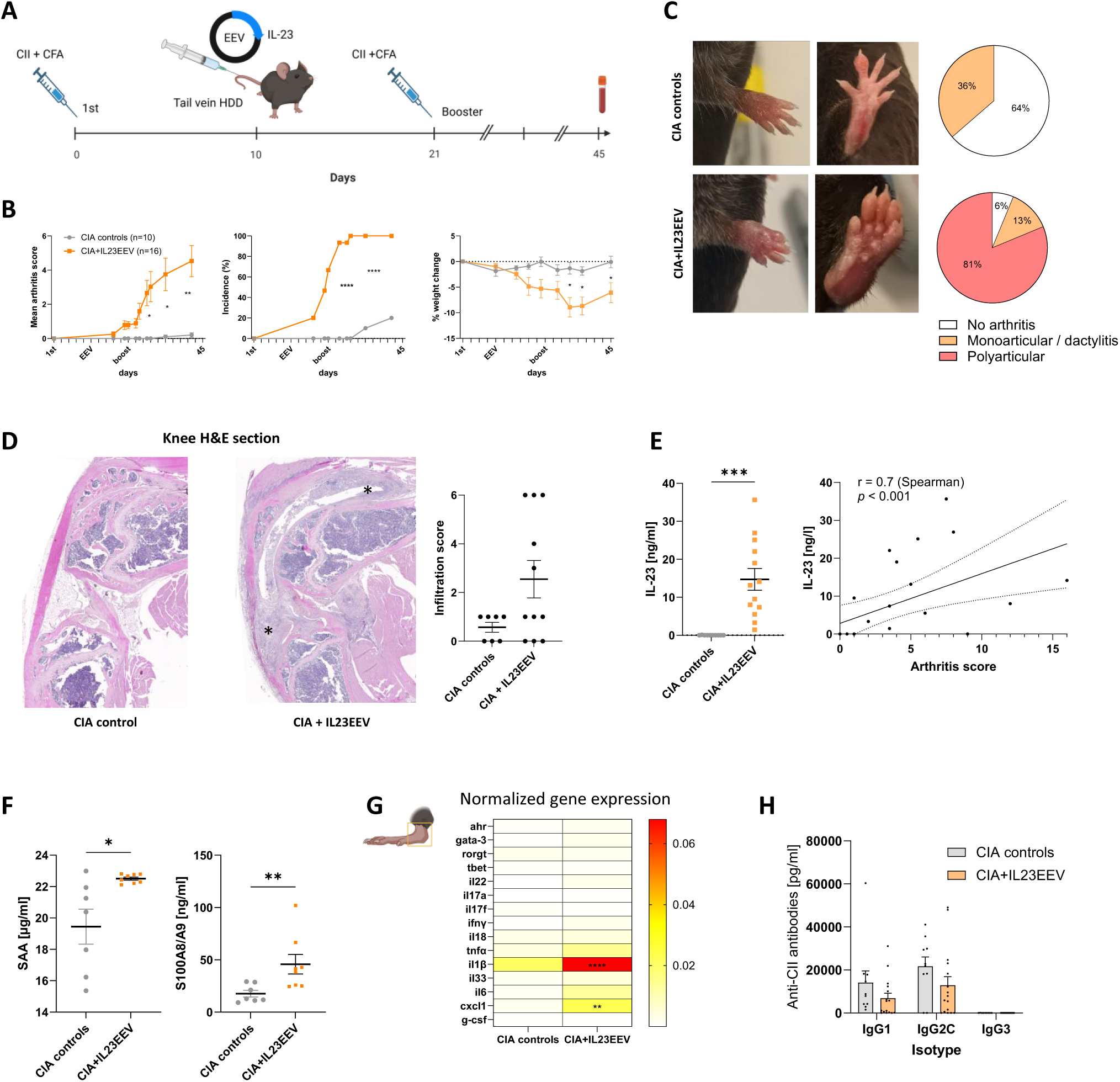
IL-23 overexpression unleashes arthritis resistance in CIA C57BL/6 mice. **(A)** Experimental design. **(B)** Clinical arthritis score (mean ± SEM), with cumulative incidence of arthritis and weight evolution in CIA mice treated with IL-23EEV (orange) compared to control (gray). **(C)** Representative photographs of paws in controls animals (upper panel) and IL-23 EEV–treated mice (lower panel), illustrating marked polyarticular inflammation, along with the distribution of affected joints. **(D)** Representative H&E-stained histological sections of knee joints showing increased synovial infiltration in the IL-23 group compared with controls. Asterisks indicate areas of synovial hypertrophy and immune cell infiltration within the synovial tissue. **(E)** Circulating plasma levels of IL-23 at day 45, with positive correlation between plasma IL-23 levels and clinical arthritis scores. **(F)** Plasma levels of serum amyloid A (SAA) and S100A8/A9. **(G)** Gene expression profiles of selected inflammatory mediators in joint tissues (ankles) by qPCR. **(H)** Humoral immune response, assessed by plasma levels of anti–type II collagen (CII) antibodies. **Statistics:** Two-way ANOVA for repeated measures and unpaired t-test for group comparisons. Statistical significance is indicated by asterisks: \**p* < 0.05, \*\**p* < 0.01, \*\*\**p* < 0.001.

**Figure 2.**
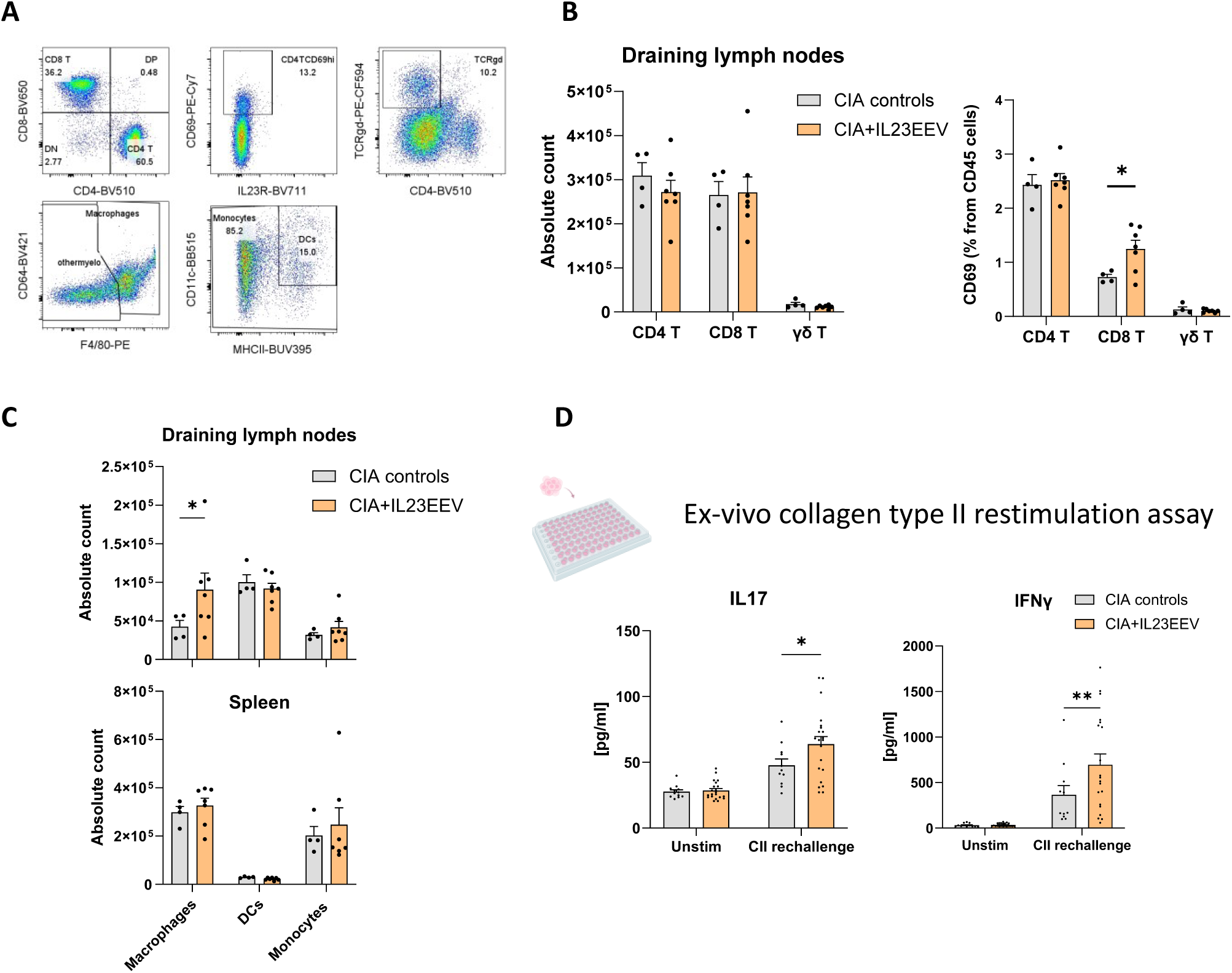
IL-23 overexpression increases specific T cells responses. **(A)** Representative flow cytometry gating strategy used to identify conventional T cells (CD19-TCRβ+CD4+ or CD8+) and γδ T cells (CD19-TCRβ-TCRγδ+), CD69-expressing T cells, macrophages (CD11b+CD64+F4/80+), monocytes (CD11b+CD64-F4/80-), and dendritic cells (CD11b+CD11c+MHCII+). **(B)** Total number of CD4+ and CD8+ conventional T cells, and γδ-T cells in the spleen and draining lymph nodes from CIA controls mice versus CIA+IL23EEV mice. **(C)** Total number of myeloid cells in the spleen and draining lymph nodes from CIA controls mice versus CIA+IL23EEV mice. **(D)** At day 45 post-immunization, splenocytes and popliteal together with inguinal lymph nodes were isolated and cells were set up in vitro for restimulation with collagen type II (100 μg/mL) or control medium. Bar plots representing quantification of IL17 and IFNγ measured by ELISA in the supernatants of cultures conditions. **Statistics:** Two-way ANOVA for repeated measures and unpaired t-test for group comparisons. Statistical significance is indicated by asterisks: \**p* < 0.05, \*\**p* < 0.01.

Histopathological examination confirmed extensive synovial and periarticular inflammatory infiltrates in the knee joints of IL-23EEV-treated CIA mice, contrasting with the minimal inflammatory changes observed in control CIA animals (Figure 1D). Circulating IL-23 concentrations were markedly increased following vector administration and positively correlated with arthritis severity (r = 0.7, p < 0.001, Figure 1E). IL23EEV treatment was also associated with elevated systemic inflammatory biomarkers, including serum amyloid A (SAA) and S100A8/S100A9 (Figure 1F). Consistent with these findings, transcriptional analysis of whole ankle tissue revealed marked upregulation of genes associated with innate immune activation and granulocyte recruitment, including *Il1b*, and *Cxcl1* mRNA (Figure 1G). Notably, disease exacerbation occurred in the absence of detectable differences in the magnitude of the antigen-specific humoral response. Levels of CII-specific IgG1, IgG2c, and IgG3 were comparable between groups, indicating that IL-23 did not enhance arthritis through increased antibody production or altered class switching (Figure 1H). Together, these results demonstrate that sustained IL-23 overexpression is sufficient to convert a poorly susceptible CIA background into a highly penetrant inflammatory arthritis model.

### IL-23 promotes antigen-specific T-cell responses and expansion of the myeloid compartment in collagen-induced arthritis

As T cells represent a major cellular target of IL-23 signaling, we next investigated the immunological mechanisms underlying the enhanced arthritis in IL-23EEV-treated CIA mice. We observed IL-23 overexpression did not significantly alter the frequencies or absolute numbers of conventional T cells or γδ T cells in either the spleen or draining lymph nodes. However, a higher proportion of CD69-expressing CD8+ T cells was detected in the draining lymph nodes of IL-23OE mice (Figure 2A-B, and supplementary figure 2). As CD69 is a well-established marker of recent T-cell activation, this finding indicates increased activation of CD8+ T cells within the draining lymph nodes. In parallel, IL-23EEV mice displayed an increased number of F4/80+CD64+ macrophages in draining lymph nodes (and frequency in the spleen), suggesting activation of the myeloid compartment (Figure 2C). Consistent with activation of the myeloid compartment, flow cytometric analysis further revealed that IL-23R expression was predominantly detected on splenic macrophages, with the highest expression observed in the MHCII^hi^CCR2^hi^ subset, in which 30.2 ± 2.6% of cells expressed IL-23R (mean ± SEM). The proportion of IL-23R-expressing cells was comparable between IL23EEV-treated and control mice (supplementary figure 3), indicating that IL-23 overexpression enhanced macrophage responses without altering receptor expression.

Next, to assess antigen-specific cellular responses, lymph node cells were cultured with splenocytes and restimulated *ex vivo* with CII. Cultures derived from IL-23EEV treated mice produced significantly higher levels of IL-17A and IFN-γ compared with lymph node cells from mice injected with control plasmids (Figure 2D). Together, these findings indicate that IL-23 promotes arthritis primarily through amplification of antigen-specific cellular immune responses and expansion of inflammatory myeloid populations rather than through broad alterations in lymphocyte numbers.

### IL-17A is required for IL-23-driven collagen-induced arthritis development

IL-17A is a major downstream effector of IL-23 signaling and has been implicated in the amplification of inflammatory responses. To determine its contribution to disease development in our model, mice were treated with a neutralizing anti-IL-17A monoclonal antibody or an isotype control (Figure 3A). IL-17A neutralization almost completely prevented arthritis development, with only occasional cases of mild mono-digital inflammation observed in treated animals. Anti-IL-17A-treated mice also exhibited significantly reduced body weight loss compared with control mice (Figure 3B-C). Consistent with macroscopic findings, histological analysis of the knee joints showed the near absence of synovial inflammatory infiltrates in anti-IL-17A-treated animals (Figure 3D). Circulating IL-23 concentrations remained unchanged between treatment groups. In contrast, systemic inflammatory markers were significantly reduced following IL-17A neutralization (Figure 3E-F). Importantly, anti-CII IgG responses were unaffected, further suggesting that protection from arthritis was not attributable to impaired antigen-specific humoral immunity (Figure 3G). Transcriptional profiling of whole ankle tissue further demonstrated a marked reduction in the expression of inflammatory mediators associated with innate immune activation, including *Il1b*, *Il6*, and *Cxcl1* mRNA (Figure 3H). Flow cytometric analysis of secondary lymphoid organs did not identify major changes in immune cell composition (data not shown). However, *ex vivo* restimulation of lymph node cells with CII demonstrated a significant reduction in IL-17A production in cultures derived from anti-IL-17A-treated mice, whereas IFN-γ secretion remained unchanged (Figure 3I). These findings suggest that IL-17A blockade selectively dampens type 17 effector responses without affecting concomitant type 1 immunity.

**Figure 3.**
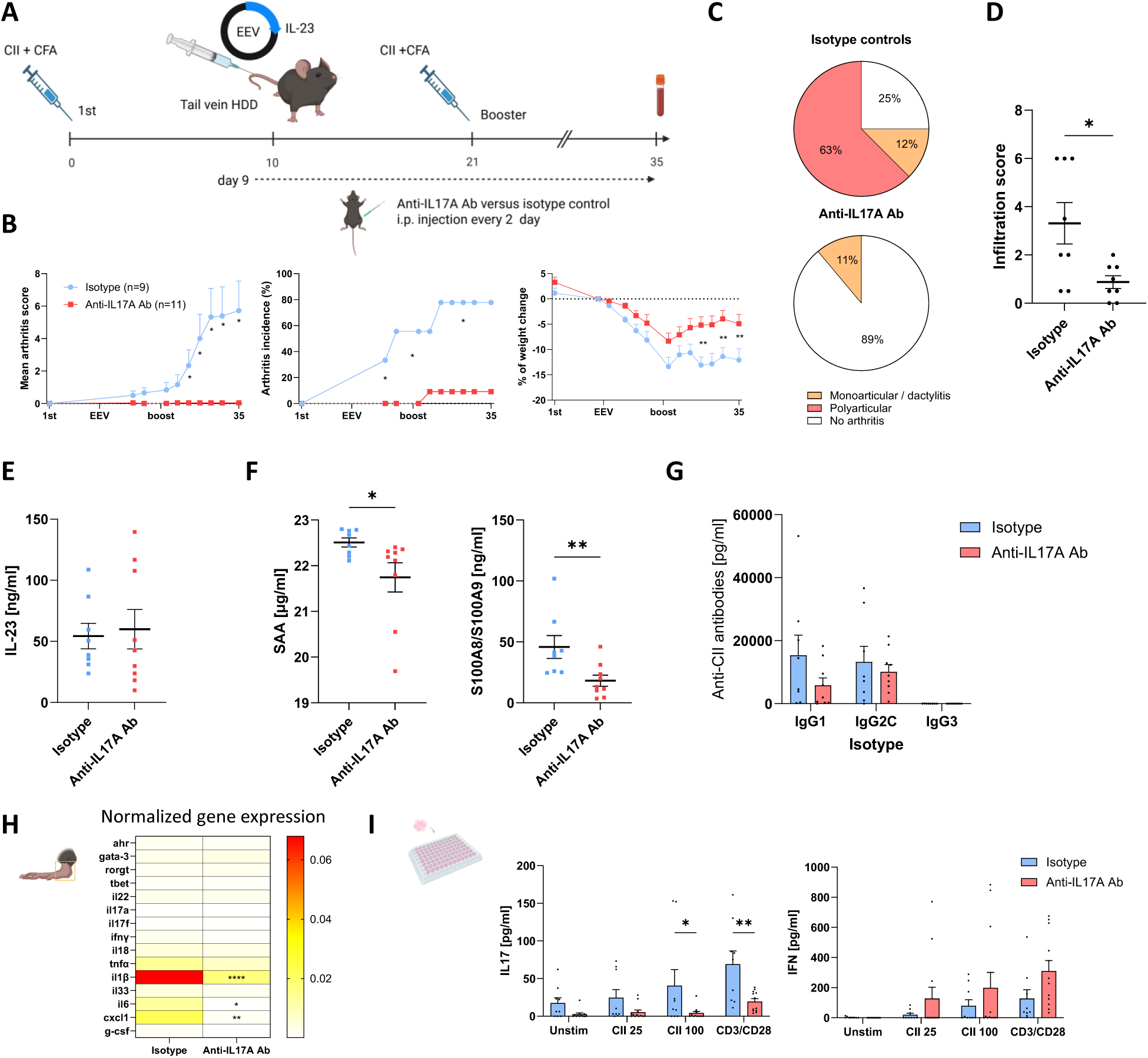
IL-17A blockade suppresses arthritis development. **(A)** Experimental design. **(B)** Clinical arthritis score (mean ± SEM) with cumulative incidence of arthritis, and weight loss in CIA+IL23EEV mice treated with anti–IL-17A (red) compared with isotype controls (blue). **(C)** Pie chart depicting the distribution of arthritis phenotype. **(D)** Infiltration score of the knee section (by H&E staining). **(E)** Circulating plasma levels of IL-23 ; **(F)** SAA and S100A8/A9 at day 35 **(G)** Plasma levels of anti–type II collagen (CII) antibodies at day 35. **(H)** Expression profiles of cytokines and inflammatory mediators in joint tissues by qPCR (ankles). **(I)** Bar plots representing quantification of IL17 and IFNγ measured by ELISA in the supernatants of draining lymph nodes after restimulation with CII. **Statistics:** Two-way ANOVA for repeated measures and unpaired t-test for group comparisons. Statistical significance is indicated by asterisks: \**p* < 0.05, \*\**p* < 0.01, *-\**p* < 0.001, \*\*\*\**p* < 0.0001.

### IL-23 pathway-related gene expression in individuals at risk of RA and early RA synovium

To investigate the relevance of the IL-23 pathway during the preclinical stages of RA, we analyzed the expression of IL-23 pathway-related genes in whole-blood transcriptomic data from 378 first-degree relatives (FDRs) of patients with RA enrolled in the SCREEN-RA cohort (1071 samples). Participants characteristics at baseline (at inclusion with blood sampling available) are presented in Table 1. Gene of interest included *Il23R, RORC, Il23A, IL12b, IL17A, IL17F, IL22, STAT3, CCR2, CCR6*, and *KLRB1*, selected based on relevant publications. Across risk groups at baseline, no significant overall group effect was observed after accounting for gene-specific differences (Wald χ² = 3.0, df = 4, p = 0.524), although a significant group-by-gene interaction was detected (Wald χ² = 51.0, df = 36, p = 0.046). To investigate the temporal dynamics of IL-23 pathway-related gene expression before RA onset, we analyzed the longitudinal data from at-risk individuals who ultimately developed RA (Figure 4A). Of 387 participants at risk at inclusion, 378 contributed to the time-varying Cox models, of whom 30 developed RA (median follow-up 3 years, CI 95% 0.8-5.3). In single-gene models adjusted for age, sex, ACPA and RF, lower expression of *KLRB1* and *RORC* was associated with a higher risk of progression (HR per SD higher expression 0.51, 95% CI 0.38–0.68, and 0.47, 0.32–0.68), and both remained significant after FDR correction (q < 0.001 and q = 0.001), Table 2. Nominal associations for *IL23R* and *IL23A* did not withstand correction (q ≥ 0.17). When all genes were considered jointly, stepwise selection retained *KLRB1* (HR 0.62, 0.43–0.90) and *RORC* (0.62, 0.38–1.02). Associations did not differ by serostatus (all q for interaction ≥ 0.66): *KLRB1* showed similar HRs in seronegative (0.56, 0.36–0.87) and seropositive individuals (0.50, 0.35–0.72). In longitudinal mixed models, *KLRB1* and *RORC* expression was lower in progressors at diagnosis than in non-progressors at last follow-up (−0.29 and −0.26 VST units; q = 0.16 for both). *RORC* tended to decline more steeply in progressors before diagnosis (slope difference −0.038 VST/year, 95% CI −0.064 to −0.012; p = 0.004, q = 0.09), whereas *KLRB1* slopes did not differ between groups (difference 0.000). Results were unchanged after additional adjustment for ACPA and RF.

**Figure 4.**
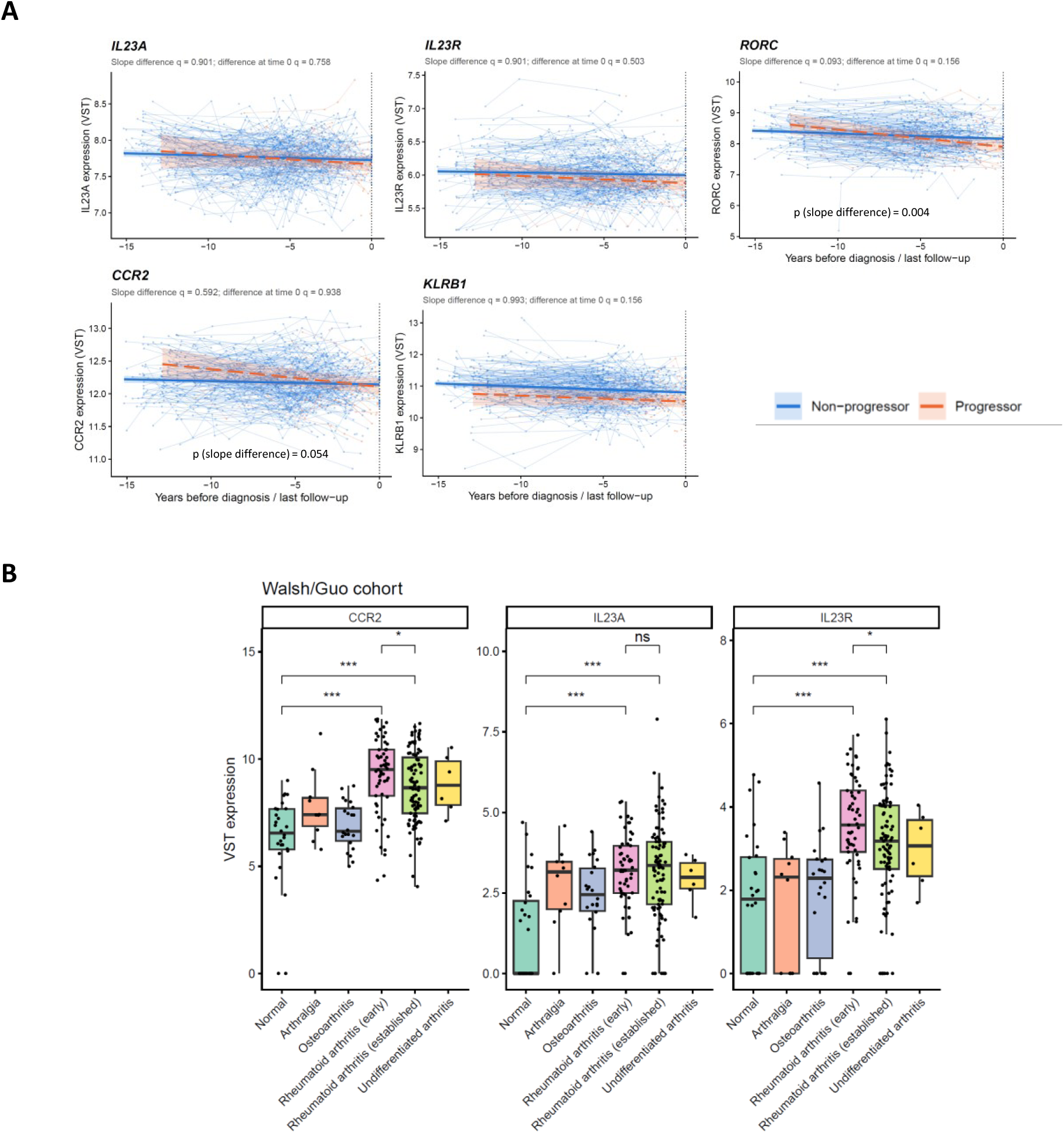
IL-23 pathway-related genes in circulation of at-risk individuals and in early rheumatoid arthritis synovium. **(A)** Longitudinal whole-blood expression of IL23A, IL23R, RORC, CCR2, and KLRB1 in individuals who developed RA (progressors, orange) and those who did not (non-progressors, blue). Thin lines represent individual expression trajectories, with points indicating individual samples. Thick lines represent fitted trends, with 95% confidence intervals. Differences in expression at the reference time point and in slopes between groups were assessed using longitudinal mixed-effects models; corresponding FDR-adjusted q-values are shown above each panel. Nominal p-values for slope differences are indicated where applicable, whereas no significant differences in expression trajectories were observed for the other genes. **(B)** Heatmap showing relative expression changes of IL-23 signature genes in each study group compared with Group 1 (controls). Color intensity reflects the magnitude and direction of differential expression relative to controls. **(C)** Box plots representing the median with interquartile range (individual samples are shown as dots) of the VST RNA-sequencing expression values for CCR2, IL23A, and IL23R in synovial biopsies from the Walsh/Guo cohort (GSE89408). Statistic : p-values between groups were estimated using a Wilcoxon rank-sum test (\**p* < 0.05, \*\**p* < 0.01, \*\*\**p* < 0.001)

**Table 1.** Baseline characteristics of SCREEN-RA participants. Demographic, clinical, and serological characteristics of the 398 participants included in the SCREEN-RA cohort, stratified according to risk group. Data are presented as numbers *(%)* or mean (SD). Group definitions are described in the Methods section. Abbreviations : ACPA, anti-citrullinated protein antibodies; BMI, body mass index; CSA, clinically suspect arthralgia; H, high-positive autoantibody level (>3× upper limit of normal); L, low-positive autoantibody level (≤3× upper limit of normal); N, negative; RA, rheumatoid arthritis; RF, rheumatoid factor; SD, standard deviation; SE, shared epitope.

|  | all | 1 | 2 | 3 | 4 | 5 | 6 |
|---|---|---|---|---|---|---|---|
| **N =** | 398 | 118 | 71 | 128 | 28 | 42 | 11 |
| **Sex (%)**, female | 318 (79.9) | 95 (80.5) | 57 (80.3) | 96 (75.0) | 24 (85.7) | 37 (88.1) | 9 (81.8) |
| **age** (mean±SD) | 50.0 (13.8) | 50.9 (12.7) | 48.4 (12.9) | 48.0 (15.1) | 53.9 (11.6) | 52.9 (13.2) | 52.6 (17.9) |
| **bmi** (mean±SD) | 24.6 (4.6) | 24.7 (4.0) | 24.1 (4.2) | 24.2 (4.6) | 24.7 (5.3) | 26.1 (6.2) | 26.2 (4.5) |
| **Smoking status** |  |  |  |  |  |  |  |
| Current (%) | 71 (18.2) | 16 (13.6) | 14 (20.0) | 18 (14.3) | 9 (32.1) | 12 (29.3) | 2 (25.0) |
| Former (%) | 125 (32.0) | 38 (32.2) | 23 (32.9) | 43 (34.1) | 8 (28.6) | 11 (26.8) | 2 (25.0) |
| Never (%) | 195 (49.9) | 64 (54.2) | 33 (47.1) | 65 (51.6) | 11 (39.3) | 18 (43.9) | 4 (50.0) |
| **Shared epitope** |  |  |  |  |  |  |  |
| absent (%) | 160 (40.5) | 55 (46.6) | 0 (0.0) | 62 (48.8) | 16 (57.1) | 23 (57.5) | 4 (36.4) |
| 1 copy (%) | 147 (37.2) | 63 (53.4) | 0 (0.0) | 52 (40.9) | 12 (42.9) | 14 (35.0) | 6 (54.5) |
| 2 copies (%) | 88 (22.3) | 0 (0.0) | 71 (100.0) | 13 (10.2) | 0 (0.0) | 3 (7.5) | 1 (9.1) |
| **ACPA** |  |  |  |  |  |  |  |
| absent (%) | 325 (81.7) | 118 (100.0) | 71 (100.0) | 77 (60.2) | 28 (100.0) | 26 (61.9) | 5 (45.5) |
| Low (%) | 43 (10.8) | 0 (0.0) | 0 (0.0) | 33 (25.8) | 0 (0.0) | 9 (21.4) | 1 (9.1) |
| Hight (%) | 30 (7.5) | 0 (0.0) | 0 (0.0) | 18 (14.1) | 0 (0.0) | 7 (16.7) | 5 (45.5) |
| **Rheumatoid factor** |  |  |  |  |  |  |  |
| absent (%) | 237 (59.5) | 88 (74.6) | 62 (87.3) | 43 (33.6) | 28 (100.0) | 13 (31.0) | 3 (27.3) |
| Low (%) | 59 (14.8) | 30 (25.4) | 9 (12.7) | 3 (2.3) | 0 (0.0) | 14 (33.3) | 3 (27.3) |
| Hight (%) | 102 (25.6) | 0 (0.0) | 0 (0.0) | 82 (64.1) | 0 (0.0) | 15 (35.7) | 5 (45.5) |
| **Ra33** |  |  |  |  |  |  |  |
| absent (%) | 210 (73.7) | 53 (75.7) | 48 (78.7) | 72 (70.6) | 20 (95.2) | 16 (55.2) | 1 (50.0) |
| Low (%) | 67 (23.5) | 17 (24.3) | 13 (21.3) | 25 (24.5) | 0 (0.0) | 12 (41.4) | 0 (0.0) |
| Hight (%) | 8 (2.8) | 0 (0.0) | 0 (0.0) | 5 (4.9) | 1 (4.8) | 1 (3.4) | 1 (50.0) |
| **CSA score** |  |  |  |  |  |  |  |
| Positive score (%) | 77 (19.4) | 0 (0.0) | 0 (0.0) | 0 (0.0) | 28 (100.0) | 42 (100.0) | 7 (70.0) |
| score (mean±SD) | 1.9 (1.3) | 1.5 (0.7) | 1.2 (0.5) | 1.2 (0.4) | 4.1 (0.8) | 4.1 (1.0) | 3.3 (1.3) |
| **SAMPLES** per patients (#, mean±SD) | 2.8 (1.5) | 3.3 (1.3) | 3.1 (1.7) | 2.7 (1.5) | 1.7 (1.1) | 2.2 (1.4) | 1.1 (0.3) |
| **FOLLOW-UP** (years, mean±SD) | 9.4 (4.6) | 10.4 (3.8) | 10.0 (4.8) | 9.8 (4.4) | 8.1 (4.5) | 7.4 (5.0) | 1.5 (3.3) |

**Table 2.** Genes associated with progression to RA in a multivariable model. Time-varying Cox regression in 378 at-risk participants (30 incident RA). Genes were selected from the 22-gene panel by stepwise selection; age, sex, ACPA and RF were included in all models. HR per 1 SD higher gene expression. p-values are not corrected for selection. ACPA, anti-citrullinated protein antibodies; CI, confidence interval; HR, hazard ratio; RF, rheumatoid factor.

| **Variable** | **HR (95% CI)** | **p** |
|---|---|---|
| **KLRB1** | 0.62 (0.43–0.90) | 0.013 |
| **RORC** | 0.62 (0.38–1.02) | 0.061 |
| **ACPA positive** | 4.18 (1.93–9.08) | <0.001 |
| **RF positive** | 2.92 (1.38–6.19) | 0.005 |
| **Age (per year)** | 0.96 (0.94–0.99) | 0.007 |
| **Female sex (ref: male)** | 2.28 (0.68–7.66) | 0.184 |

Finally, to verify whether IL-23 pathway activation was also detectable at the site of disease, we analyzed a publicly available RNA-sequencing dataset of synovial biopsies from patients spanning the spectrum of inflammatory arthritis (GSE89408)^25,26^. Synovial expression of IL23R, IL23A, and CCR2 was significantly increased in RA compared with healthy synovium and non-RA controls (Figure 4B). Remarkably, IL23R expression was most pronounced in early RA compared with established disease, suggesting that IL-23-responsive immune populations may be particularly enriched during disease initiation. Together, these cross-sectional and longitudinal analyses identify differences in IL-23 pathway-related gene expression before clinical RA onset and in synovial tissue during early disease.

## Discussion

The mechanisms governing the transition from systemic autoimmunity to arthritis remain incompletely understood in RA. In the present study, we provide complementary experimental and translational evidence implicating IL-23 in the transition from systemic autoimmunity to inflammatory arthritis. Sustained IL-23 overexpression converted an otherwise poorly susceptible CIA model into a highly penetrant inflammatory arthritis characterized by synovial inflammation, activation of innate immune pathways, and enhanced antigen-specific T-cell responses. Importantly, these effects occurred without alterations in collagen-specific antibody production, supporting a role for IL-23 in amplifying inflammatory effector mechanisms in the presence of established antigen-specific autoimmunity.

Our findings provide experimental support for a multi-step model of RA development in which systemic autoimmunity alone is insufficient to induce arthritis. This concept is highly relevant to human RA, where ACPA positivity alone confers risk but does not inevitably lead to disease^27^. Collectively, our data suggests that IL-23 could act as a molecular switch capable of converting pre-existing autoimmunity into tissue inflammation. Our findings complement those reported by Pfeifle et al., who demonstrated that IL-23 promotes the transition from autoimmunity to arthritis through modulation of autoantibody pathogenicity via altered Fc glycosylation^16^. In that study, IL-23 reprogrammed Th17 cells to suppress St6gal1 expression in differentiating antibody-producing cells through mechanisms dependent on IL-21 and IL-22, resulting in the generation of more arthritogenic autoantibodies. In our model, however, the arthritogenic effects of IL-23 were largely dependent on its downstream effector cytokine IL-17A. Mechanistically, our data suggest that IL-23 promotes the expansion and activation of pathogenic IL-17A-producing T cells, which subsequently orchestrate a potent granulocytic inflammatory response within the joint. Consistent with this hypothesis, arthritic joints displayed increased expression of innate inflammatory mediators involved in neutrophil recruitment and activation, including *Il1b*, and *Cxcl1*. These changes were also accompanied by elevated circulating levels of S100A8/A9, a biomarker of neutrophil activation and disease activity in RA, which correlated with arthritis score in our model (supplementary figure 4)^28^.

Notably, IL-23R expression was highest on MHCII^hi^CCR2^hi^ macrophages, a subset associated with inflammatory monocyte recruitment and antigen presentation^29^. CCR2-dependent recruitment of immune cells to inflammation sites has been implicated in the development of synovitis^30–32^. Although our findings do not establish a direct functional role for IL-23R-expressing macrophages in arthritis development, they raise the possibility that IL-23 may also modulate inflammatory myeloid responses. Hence, IL-23 may promote disease transition not only through pathogenic T-cell responses but also through direct amplification of inflammatory myeloid circuits. Altogether, these findings suggest that IL-23 acts through non-mutually exclusive mechanisms, both on the pathogenicity of autoantibodies and on the activation of IL-17-driven cellular and myeloid inflammatory pathways^33,34^.

Strikingly, neutralization of IL-17A almost completely prevented arthritis development, abolished synovial inflammation, and suppressed the expression of key inflammatory mediators in joint tissue. These results demonstrate that early articular manifestations in this model are highly dependent on the IL-23/IL-17A axis. In RA synovium, IL-23 and IL-17A-producing cells have been shown to colocalize within inflamed tissue, with their abundance demonstrating a positive correlation across samples^35^. A potential concern is that the prominent dependence of disease on the IL-23/IL-17 axis in our model may resemble mechanisms more commonly associated with psoriatic arthritis than established RA. Indeed, therapeutic inhibition of both IL-17 and IL-23 has shown substantial efficacy in psoriatic arthritis but has failed to demonstrate meaningful clinical benefit in established RA^36,37^. However, our model was developed in the context of antigen-specific collagen-induced autoimmunity and required prior immunization for arthritis development, as IL-23 overexpression alone was insufficient to induce joint disease.

Moreover, accumulating evidence suggests that this pathway may be particularly relevant during the earliest phases of disease and several observations support the relevance of these findings to human disease^14,38^. A recent single cell study identified a population of CCR2+ CD4+ T cells enriched in individuals at risk of RA that express high levels of *IL23R* together with multiple components of a type 17 transcriptional program, including *CCR6*, *KLRB1*, *IL22*, *IL17A*, and *TNF*^39^. Interestingly, this subset was specifically associated with ARI seropositivity. The observed decline in RORC and KLRB1 expression in peripheral blood before RA onset may reflect changes in circulating immune-cell populations, potentially including the recruitment of these cells into peripheral tissues such as the synovium as suggested by others^40^. However, this hypothesis remains speculative and further studies are needed to determine whether immune-cell migration contributes to these longitudinal expression changes. Interestingly, analysis of an independent synovial transcriptomic dataset demonstrated increased expression of *IL23R*, *IL23A*, and *CCR2* in RA synovium, with *IL23R* expression being most pronounced in early disease. I*L23R* expression is largely confined to activated type 17 lymphocyte populations and is induced by pro-inflammatory cytokines such as IL-6, IL-21, and IL-1β through STAT3-dependent pathways^41^. Thus, increased *IL23R* expression may reflect the expansion or activation of IL-23-responsive immune populations during the preclinical phase of RA. These findings may help explain why therapeutic blockade of IL-23 has failed in established RA. It is consistent with genetic studies implicating functional *IL23R* variants in susceptibility to several autoimmune diseases, including inflammatory bowel disease, psoriasis, and RA^42,43^. We therefore propose that IL-23/IL17A signaling may be most important during the earliest phases of RA pathogenesis and disease transition rather than during the maintenance and perpetuation of established chronic synovitis.

These observations have important implications for disease interception strategies. Recent observational studies in psoriasis have reported a lower incidence of psoriatic arthritis among patients treated with IL-23 inhibitors compared with TNF or IL-17 inhibitors^44^, and prospective clinical trials are currently evaluating whether IL-23 blockade can prevent disease onset in individuals at risk of psoriatic arthritis^45^. In RA, the strongest evidence for disease prevention has emerged from interventions targeting T-cell activation, notably abatacept in the APIPPRA and ARIAA trials^46,47^. Taken together, our experimental and translational findings support a model in which IL-23-responsive immune populations contribute to disease transition, raising the possibility that therapeutic targeting of the IL-23 pathway could represent also a rational strategy for preventing progression from systemic autoimmunity to clinically manifest RA.

Several limitations should be acknowledged. The study relies on forced IL-23 overexpression and collagen immunization, which may not fully recapitulate the complexity of human disease. Furthermore, the translational observations are cross-sectional and do not establish a causal relationship between *IL23R* expression and progression to inflammatory arthritis. Although the longitudinal analyses provide information on gene-expression trajectories before RA onset, the limited number of progressors and the correlation between gene-expression measures constrain the conclusions that can be drawn about independent associations with disease progression. Future studies integrating longitudinal at-risk cohorts, synovial tissue analyses, and single-cell approaches will be required to determine whether IL-23-responsive cellular populations directly predict disease transition in humans.

In conclusion, our findings identify IL-23 as an experimental driver of inflammatory arthritis in the context of established antigen-specific autoimmunity and implicate IL-23 pathway-related gene expression modification in the preclinical and early stages of human RA. In mice, IL-23 promoted arthritis through IL-17A-dependent mechanisms involving both adaptive and innate inflammatory responses. In humans, increased IL23R and CCR2 expression in individuals with RA-related autoimmunity and enrichment of IL-23 pathway-related genes in early RA synovium provide complementary translational evidence. Longitudinal analyses further revealed that declining RORC and KLRB1 whole blood expression was associated with a higher risk of RA onset, highlighting dynamic changes in gene expression preceding disease development.. These findings support further investigation of IL-23 as a potential target for disease prevention before the establishment of chronic synovitis.

## Acknowledgment

This work was financially supported by the De Reuter Foundation, the Jean and Linette Warnery Foundation (2025 Rheumatology Research Prize), the Jean Tua Foundation, and the Private Foundation of Geneva University Hospitals (HUG). M.J. gratefully acknowledges their generous support.

The authors thank Prof. Ioannis Xenarios and the Health 2030 Genome Center (https://www.health2030genome.ch/; Campus Biotech, Geneva, Switzerland), and acknowledge the European projects SYSCID (A Systems Medicine Approach to Chronic Inflammatory Diseases; https://syscid.eu/) and PerPrevCID (Personalised Disease Prediction and Prevention in Chronic Inflammatory Disorders ; www.perprev-cid.eu) cohorts for providing access to the data used in this study.

The authors also thank Novartis for generously providing the anti-mouse IL-17A antibodies used in this study.

## Supplementary figures

**Supplementary Figure 1.**
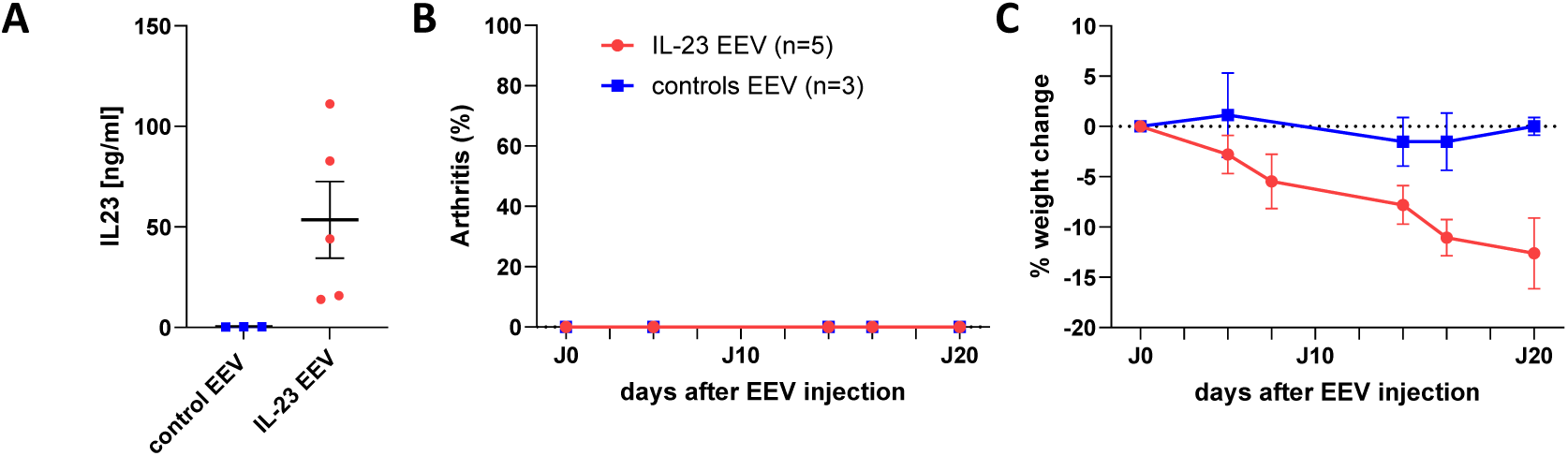
IL-23 overexpression is insufficient to induce arthritis in non-immunized C57BL/6 mice. **(A)** Serum IL-23 concentrations following administration of control EEV or IL-23EEV. **(B)** Arthritis incidence over time showing no development of clinical arthritis in non-immunized mice despite sustained IL-23 overexpression. **(C)** Percentage body weight change over time. IL23EEV induced weight loss compared with control EEV-treated mice. Data are shown as mean ± SEM.

**Supplementary figure 2.**
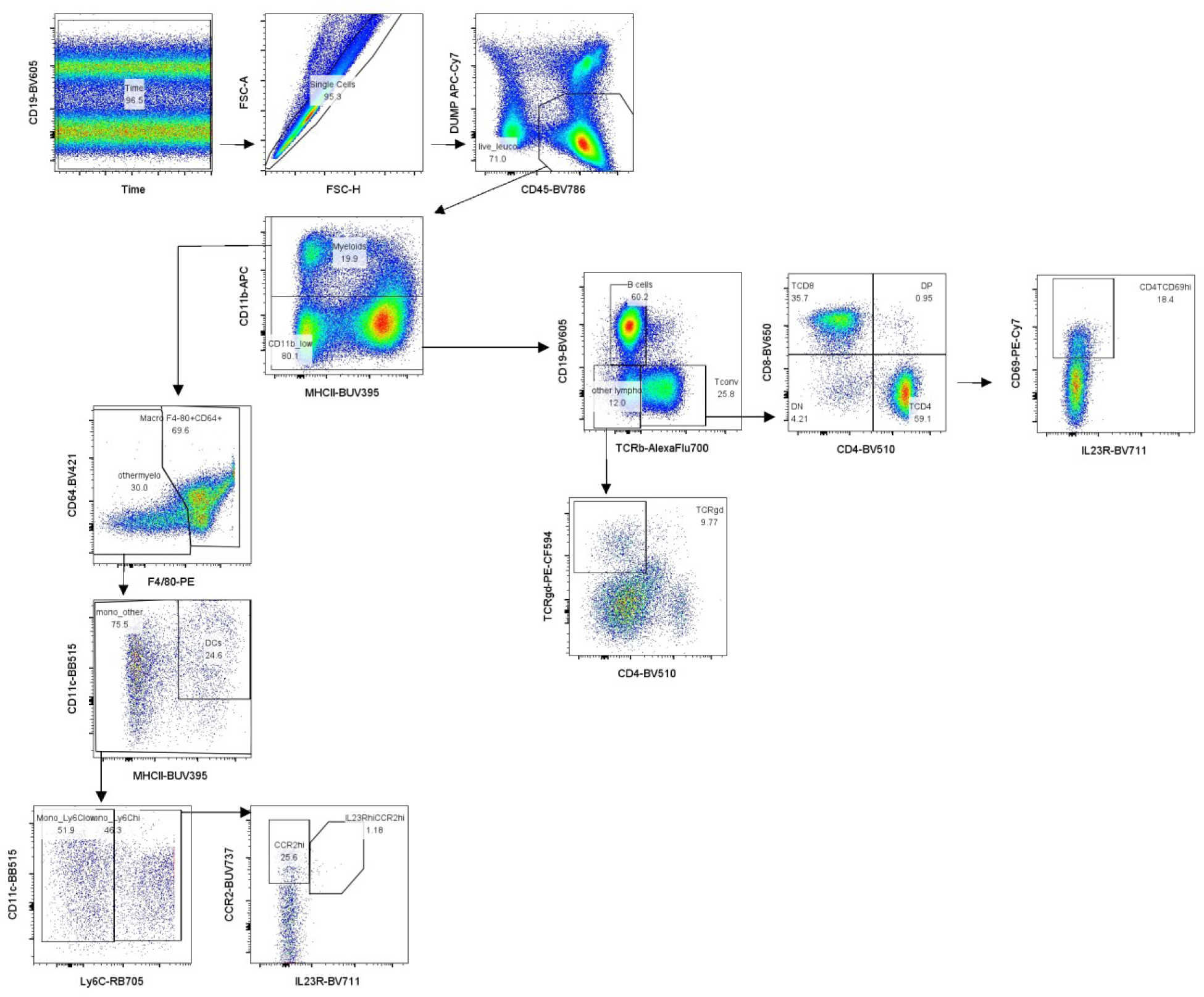
Gating strategy. Representative gating strategy used to identify the major immune cell populations analyzed in this study.

**Supplementary figure 3.**
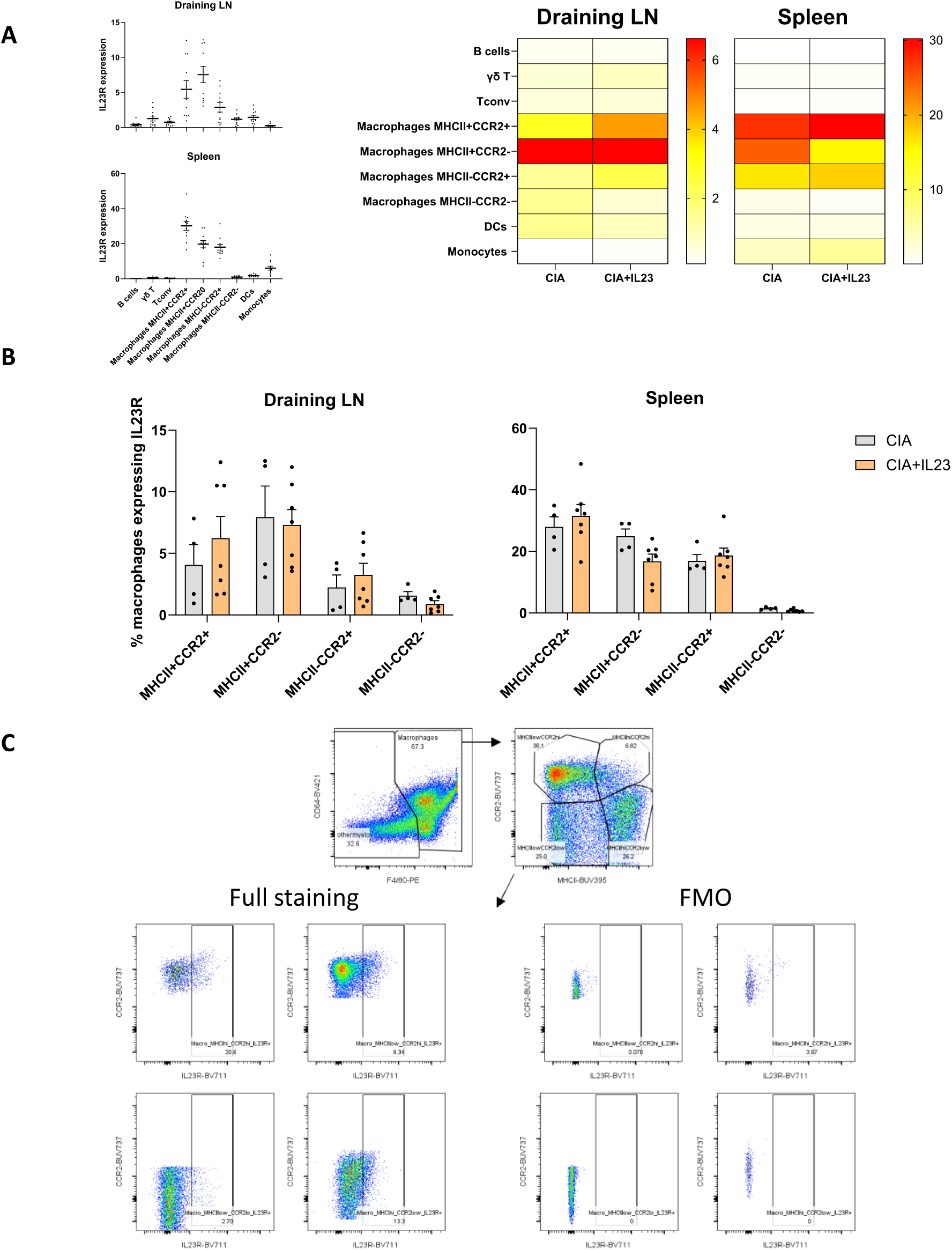
Distribution of IL-23R expression across selected immune cell populations. **(A).** Dot plots depicting the frequency of IL-23R⁺ cells within the indicated immune cell subsets in the spleen and draining lymph nodes, and heat maps showing no differences in CIA control versus IL-23EEV-treated mice, as determined by flow cytometry. Color intensity represents the mean percentage of IL-23R-expressing cells within each parent population. **(B)** Frequency of IL-23R⁺ cells among macrophage subsets from the spleen and draining lymph nodes of CIA control and IL-23EEV-treated mice. Bars represent mean ± SEM. **(C)** Representative flow cytometry plots illustrating the gating strategy used to identify IL-23R-expressing macrophage subsets, together with fluorescence-minus-one (FMO) controls used to define IL-23R positivity.

**Supplementary figure 4.**
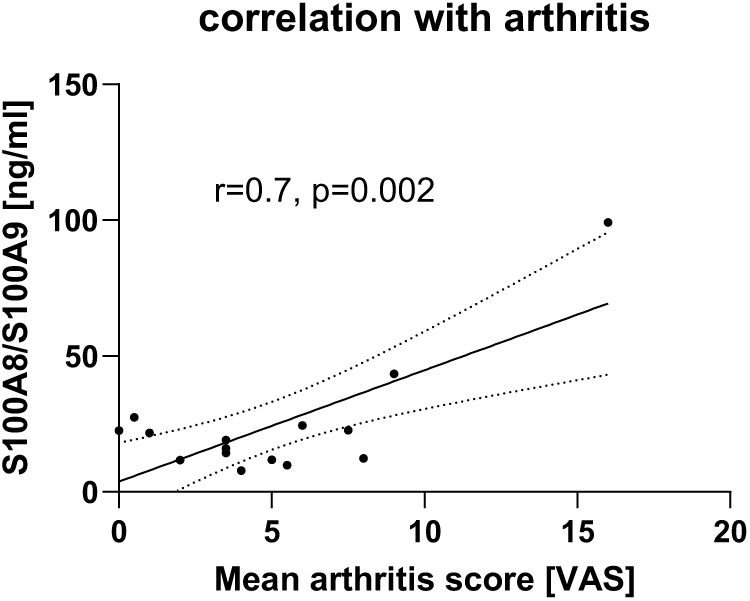
Correlation between levels of circulating calprotectin (S100A8/S100A9) and visual arthritis score. Statistical significance was assessed using Pearson’s correlation test.

## Notes

https://www.ncbi.nlm.nih.gov/geo/query/acc.cgi?acc=GSE89408

### Competing Interest Statement

The authors have declared no competing interest.

