## Supplementary methods for "IL-23 licenses the transition from preclinical autoimmunity to inflammatory arthritis through IL-17A-dependent mechanisms"

Jarlborg M., et al.

**Supplementary methods**

Plasma biomarkers and antibody measurements

Plasma concentrations of IL-23 were quantified using the Mouse IL-23 DuoSet ELISA kit (R&D Systems, Bio-Techne, Minneapolis, MN, USA, Cat. DY1887). Systemic inflammation was assessed by measuring serum amyloid A (SAA) and S100A8/S100A9 heterodimer (calprotectin) using DuoSet ELISA kits (R&D Systems, Bio-Techne, Minneapolis, MN, USA), according to the manufacturer's instructions. Collagen-specific humoral responses were evaluated by measuring anti-type II collagen (CII) IgG1, IgG2c, and IgG3 antibodies using commercially available ELISA kits (Chondrex, Redmond, WA, USA, Cat. 20311, 20315 & 20314 ).

*Ex vivo* antigen-specific restimulation assays

Single-cell suspensions from spleens and draining lymph nodes (popliteal and inguinal) were prepared at sacrifice and cultured in 96-well U-bottom plates at a density of 2 × 10^5 cells per well. Cells were cultured in RPMI-1640 supplemented with 10% fetal bovine serum, 1% penicillin-streptomycin, 2 mM L-glutamine, and 50 μM β-mercaptoethanol. Cells were restimulated with chicken type II collagen (100 ng/mL) for 24 h at 37°C in a humidified incubator containing 5% CO₂. Control conditions included unstimulated cells cultured in medium alone and polyclonal stimulation using Dynabeads™ Mouse T-Activator CD3/CD28 (Thermo Fisher Scientific, Waltham, MA, USA), according to the manufacturer's instructions. Following incubation, culture supernatants were collected after centrifugation and stored at −80°C until analysis. Cytokine concentrations in the cultured supernatant were quantified by ELISA, including IL-17A (Mouse IL-17 DuoSet ELISA, R&D Systems, Bio-Techne; Cat. DY421) and interferon-γ (Mouse IFN-γ Uncoated ELISA Kit, Invitrogen, Carlsbad, CA, USA, Cat. 88-7314-88), according to the manufacturers' protocols.

Histology

Knee joints were dissected, fixed in 4% paraformaldehyde, and decalcified in EDTA for 3 weeks prior to paraffin embedding. Serial 5-μm sections were prepared and stained with hematoxylin and eosin (H&E) for histopathological evaluation. Histological scoring was performed in a blinded manner by two independent observers using a semi-quantitative scoring system assessing synovial and skin inflammation. Briefly, inflammation was graded from 0 to 3 according to the extent of inflammatory cell infiltration (0, normal synovium; 1, mild infiltration; 2, moderate infiltration; 3, severe infiltration with dense inflammatory cell aggregates). Synovial hyperplasia was graded from 0 to 3 based on synovial lining thickness (0, 1–2 cell layers; 1, 3–4 cell layers; 2, 5–6 cell layers; 3, >6 cell layers with loss of normal architecture).

Reverse transcription (RT)-quantitative polymerase chain reaction (qPCR)

Total RNA was extracted from whole ankle joints and ear tissues following mechanical homogenization in TRIzol reagent using the RNeasy Mini Kit (Qiagen, Hilden, Germany), according to the manufacturer's instructions. RNA concentration and purity were determined spectrophotometrically, and 500 ng of total RNA was reverse-transcribed using PrimeScript™ Reverse Transcriptase (Takara Bio, Kusatsu, Japan).

Quantitative real-time PCR (qRT-PCR) was performed using SYBR Select Master Mix (Thermo Fisher Scientific) on a QuantStudio™ 6 Pro Real-Time PCR System (Thermo Fisher Scientific). Relative gene expression was calculated using the comparative Ct (2^-ΔΔCt^) method and normalized to *Gapdh* and *L32* expression. All primers were synthesized by Eurofins Genomics. Primer sequences (5′–3′) were as follows: *Ahr* (F: CAGCGGCGCCAACATCACCT; R: AGGCTGGCCAGGCGGTCTAA), *Gata3* (F: AGAACCGGCCCCTTATCAA; R: AGTTCGCGCAGGATGTCC), *Rorc* (F: CCTCTACACGGCCCTGGTT; R: GATGTTCCACTCTCCTCTTCTCTTG), *Tbx21* (T-bet) (F: CAACAACCCCTTTGCCAAAG; R: TCCCCCAAGCAGTTGACAGT), *Il22* (F: TTGACACTTGTGCGATCTCTGA; R: CGGTTGACGATGTATGGCTG), *Il17a* (F: CCACGTCACCCTGGACTCTC; R: CTCCGCATTGACACAGCG), *Ifng* (F: AGACAATCAGGCCATCAGCA; R: TGGACCTGTGGGTTGTTGAC), *Il18* (F: TCAGACAACTTTGGCCGACT; R: CAGTCTGGTCTGGGGTTCAC), *Tnf* (F: AGTTCTATGGCCCAGACCCT; R: GTCTTTGAGATCCATGCCGT), *Il1b* (F: TGTGAAATGCCACCTTTTGA; R: GTGCTCATGTCCTCATCCTG), *Il33* (F: GGTGTGGATGGGAAGAAGCTG; R: GAGGACTTTTTGTGAAGGACG), *Il6* (F: TAGTCCTTCCTACCCCAATTTCC; R: TTGGTCCTTAGCCACTCCTTC), and *Cxcl1* (F: ACTCAAGAATGGTCGCGAGG; R: GTGCCATCAGAGCAGTCTGT).

Flow cytometry

Single-cell suspensions were prepared from spleens and draining lymph nodes (popliteal and inguinal, bilaterally). Spleens were submitted to red blood cell lysis prior to staining. Cells were pelleted by centrifugation, resuspended in PBS containing 0.5% bovine serum albumin (BSA), and incubated with Fc receptor blocking reagent (Miltenyi Biotec, Bergisch Gladbach, Germany). Cells were then stained with fluorochrome-conjugated antibodies against CD45 (BV786, ref. 564225, BD Biosciences), Anti-Mouse I-A, I-E (BUV395, ref. 589244, BD Biosciences), CCR2 (BUV737, ref. 759593, BD Biosciences), CD11c (BB515, ref. 565586, BD Biosciences), Ly6C (RB705, ref. 570264, BD Biosciences), F4/80 (PE, ref. 565410, BD Biosciences), TCRγδ (PE-CF594, ref. 563532, BD Biosciences), CD69 (PE-Cy7, ref. 552879, BD Biosciences), CD11b (APC, ref. 553312, BD Biosciences), TCRβ (R718, ref. 567299, BD Biosciences), CD64 (BV421, ref. 139309, BioLegend), CD4 (BV510, clone GK1.5, ref. 569249, BD Biosciences), CD19 (BV605, ref. 115539, BioLegend), CD8 (BV650, ref. 100742, Biolegend), IL-23R (BV711, ref. 744372, BD Biosciences), and Ly6G (APC-Cy7, ref. 127624, Biolegend). Viable cells were identified using DRAQ7 viability dye (BioStatus), which was added immediately prior to acquisition. Samples were acquired on an LSRFortessa flow cytometer (BD Biosciences) and analyzed using FlowJo software version 10 (BD Life Sciences).

**Supplementary table 1.** List of genes selected a priori to characterize IL-23 pathway activity, including genes involved in IL-23 signaling, type 17 immunity, and associated cellular markers. Gene symbols and corresponding Ensembl gene identifiers used for transcriptomic analyses are shown.

| **GENE_ID** | **ENSEMBL_ID** |
| --- | --- |
| IL23R | ENSG00000162594.17 |
| IL23A | ENSG00000110944.9 |
| IL12B | ENSG00000113302.6 |
| RORC | ENSG00000143365.21 |
| IL17A | ENSG00000112115.7 |
| IL17F | ENSG00000112116.11 |
| IL22 | ENSG00000127318.11 |
| CCR6 | ENSG00000112486.19 |
| STAT3 | ENSG00000168610.18 |
| KLRB1 | ENSG00000111796.4 |
| CSF2 | ENSG00000164400.6 |
| CCR2 | ENSG00000121807.8 |
| IL1A | ENSG00000115008.7 |
| IL1RN | ENSG00000136689.20 |
| IL18 | ENSG00000150782.14 |
| IL18BP | ENSG00000137496.20 |
| IL33 | ENSG00000137033.13 |
| IL1RL1 | ENSG00000115602.18 |
| IL25 | ENSG00000166090.8 |
| HMGB1 | ENSG00000189403.16 |
| S100A8 | ENSG00000143546.11 |
| S100A9 | ENSG00000163220.12 |
| S100A7 | ENSG00000143556.10 |
| S100A12 | ENSG00000163221.9 |
| TSLP | ENSG00000145777.15 |
| HSPA1A | ENSG00000204389.11 |
| REG3G | ENSG00000143954.14 |
| LCN2 | ENSG00000148346.13 |
| PADI4 | ENSG00000159339.13 |
| PADI2 | ENSG00000117115.14 |
| IL6 | ENSG00000136244.13 |
| TNF | ENSG00000232810.5 |
| CSF3 | ENSG00000108342.14 |
| TGFB1 | ENSG00000105329.12 |
